# A semantotopic map in human hippocampus

**DOI:** 10.1101/2025.10.31.685959

**Authors:** Elizabeth A. Mickiewicz, Melissa Franch, Kalman A. Katlowitz, Ana G. Chavez, Hanlin Zhu, Assia Chericoni, Xinyuan Yan, James L. Belanger, Taha Ismail, Danika L. Paulo, Alica M. Goldman, Vaishnav Krishnan, Atul Maheshwari, Eleonora Bartoli, Sarah R. Heilbronner, Nicole R. Provenza, Sameer A. Sheth, Benjamin Y. Hayden

**Affiliations:** Neurosurgery, Baylor College of Medicine; Neurology, Baylor College of Medicine; Electrical and Computer Engineering and Neuroengineering Initiative, Rice University

## Abstract

The hippocampus plays a central role in encoding abstract conceptual and semantic information. However, little is known about the topography of that encoding. We leveraged the rare opportunity to examine neural responses densely along a small portion of the human hippocampus, specifically, the mediolateral axis of the anterior body. We collected responses to passive language listening using Neuropixels probes in three anesthetized patients during clinically indicated neurosurgical procedures. We computed semantic tuning functions for each recording site by regressing threshold crossing events and single unit responses against semantic embeddings from GPT-2, Word2Vec, and SBERT. We find that tuning functions of more distant recording sites are more dissimilar, supporting the hypothesis semantotopic organization. Multiple semantic features showed systematic changes along that axis, including animacy, concreteness, and familiarity; notably, effects were individual-specific. Surprisingly, we also found a small but significant increase in semantic similarity as a function of distance between recording sites, on a shorter spatial scale, suggesting a modest periodic organization. Together, these results demonstrate the presence of a multiscale functional organization of semantics in the hippocampus.

## INTRODUCTION

The brain encodes the meanings of words we hear using a population code that has at least some features in common with encoding in large language models (LLMs, Chavez et al., 2025; Franch et al., 2025; Jamali et al., 2024; Karkowski et al., 2025; Katlowitz et al., 2025). However, unlike large language models, neurons are densely packed into the physical space of the brain. This physicality imposes both opportunity and obligation, as costs of communicating and coordinating learning across distances may determine the organization of function (Bosman & Aboitiz, 2015; Park & Friston, 2013; Fotiadis et al., 2024; Bullmore & Sporns, 2012). These factors have led to a great deal of interest in the question of functional organization of association regions of the brain (Yeo et al., 2015; Mutschler et al., 2009), including the hippocampus (Robinson et al., 2015). Understanding this organization can help us advance theories of brain development, improve interpretation of lesion and MRI studies, drive theories of brain dysfunction, and inspire more efficient computer architectures (Fair et al., 2009; Klimes et al., 2016; Müller & Knight, 2006; Mehonic & Kenyon, 2022).

There is considerable evidence for macroscale organization of abstract information, including the existence of specialized face patches (Kanwisher et al., 1997; Tsao et al., 2008; Leopold et al., 2006), place-selective areas (Epstein et al., 1999), language-selective regions (Fedorenko et al., 2024), and, possibly, functionally homogeneous patches in inferotemporal cortex (Sato et al., 2013). Likewise, there is some evidence for a broad segregation of language so that, for example, nouns and verbs are localized to different regions (Huth et al., 2016; Binder et al., 2009; Friederici et al., 2000; Binder et al., 2005). However, it remains unclear how, or even if, that information is organized *within* regions or subregions. The majority of information about subareal functional organization (i.e., maps) comes from primary sensory and motor cortices, such as tonotopy in the auditory cortex and somatotopy in the somatosensory cortex (Gilbert & Wiesel, 1983; Linden & Schreiner 2003; Rao et al., 1995; Humphries et al., 2010; Kuljis & Rakic, 1990; Sanchez Panchuelo et al., 2018).

However, in the large segment of association cortex between sensory and motor regions, principles of functional organization are considerably less clear (Patel et al., 2014). Indeed, many studies have looked for maps relating to information coded within the association cortex and found very little. For example, there is little evidence for organization within face patches of the macaque at the millimeter level using fMRI, let alone the micrometer level examining individual neurons (Janssens et al., 2014; Finzi et al., 2021). In a similar vein, concept-sensitive neurons appear to have little detectable organization; this lack of organization has even been proposed to be part of how these neurons support flexible associations (De Falco et al., 2016). Thus, it is possible that intra-areal organization is largely unstructured. However, there is some evidence that is at least suggestive of millimeter scale functional subareal organization in the temporal lobe (Afraz et al., 2006; Moeller et al., 2017; Grill-Spector & Weiner, 2014; Leonard et al., 2024). Moreover, the question of semantic tuning has not been directly addressed, due to two difficulties of collecting dense measurements in semantically sensitive regions and in the difficulty of quantifying semantics.

This second problem can be addressed using semantic embeddings given by LLMs. LLMs such as GPT-2 provide a quantitative representation of semantics in a high-dimensional space (Radford et al., 2019). Broadly speaking, LLMs build an embedding space for semantic information in a bottom-up manner, following the principle that a word’s meaning can be defined by the other words that tend to occur nearby it. We have previously found that semantic embeddings derived from LLMs can provide a good accounting of neuronal firing rates in the hippocampus (Franch et al., 2025; Katlowitz et al., 2025A and B; Chavez et al., 2025). Semantic embeddings therefore offer the ability to quantitatively assess the high dimensional “tuning curves” for semantic coding. We can then compare these tuning curves to evaluate the spatial organization of semantic coding in the human hippocampus.

The hippocampus is especially interesting in this regard due to its putative role in conceptual representation (Mack et al., 2018; Kafkas et al., 2024; Theves et al., 2021; Constantinescu et al., 2016; Behrens et al., 2018), including for language semantics (Blank et al., 2016; Davachi, 2006; Wolna et al., 2025). We and others have argued that this role includes encoding word meanings during speech listening (Franch et al., 2025, Katlowitz et al., 2025B, Dijksterhuis et al., 2024, Sassenhagen & Fiebach, 2020; Fedorenko et al., 2018). There is plentiful evidence for functional organization within the hippocampus. For example, along the rostrocaudal axis, there is evidence for a relative selectivity for emotion and stress regulation in more rostral regions and cognitive function in more caudal regions (Fanselow & Dong, 2010). And, between subfields, there appears to be specialization for processes like pattern completion/ separation (Yassa and Stark, 2011). However, the question of semantotopy remains unanswered. Here we leveraged the rare opportunity to examine encoding of linguistic features along a single axis (the mediolateral axis) of the hippocampus using Neuropixels probes (Steinmetz et al., 2018).

## RESULTS

We performed intraoperative hippocampal recordings with Neuropixels probes in three patients undergoing anterior temporal lobectomy for drug resistant epilepsy (Katlowitz et al., 2025A). Hippocampal recordings were conducted in the anterior body after resection of the lateral temporal cortex and prior to resection of the mesial temporal structures (**Figure 1A**).

**Figure 1.**
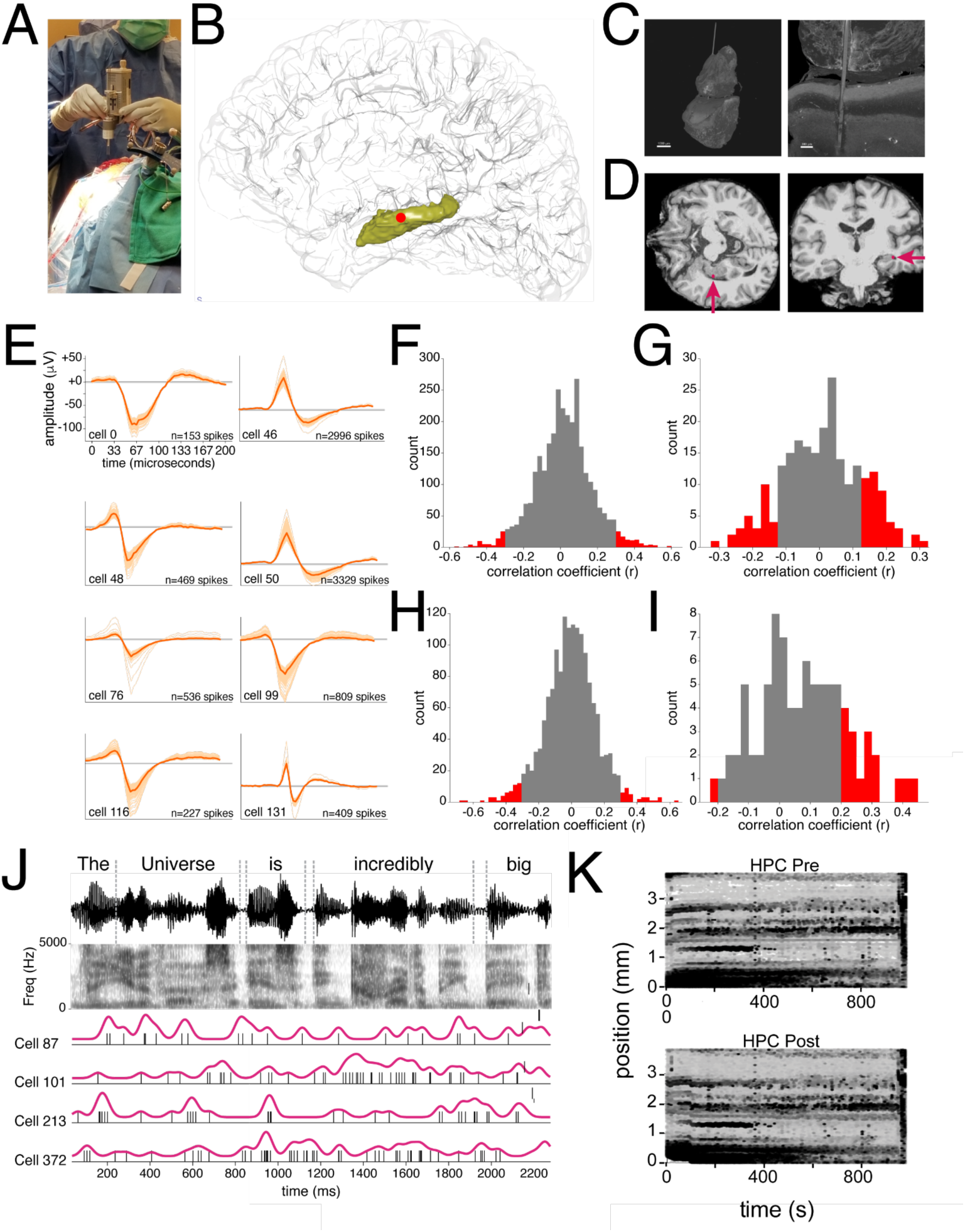
Intraoperative Neuropixels recordings in the human hippocampus during word listening task. **A.** Photograph of OR set-up. **B.** Probe entry site for patients warped onto a canonical brain, illustrated with a red dot within the hippocampus, shown in yellow. **C.** In one patient (patient 2), we managed to extract the probe along with the tissue. 3D reconstruction of microCT identifying the probe within resected hippocampal tissue (left) with a coronal slice identifying the probe penetrating the hippocampus (right). **D.** Axial (left) and coronal (right) sections of a T1 MRI for patient 2. **E.** Example units from patient 3. **F.** Continuous spiking activity from four example neurons from patient 2 during listening. Timestamped words shown in relation to the audio waveform and spectrogram. **G.** Drift maps illustrating the low drift rates and high stability of Neuropixel recordings in the human hippocampus. X-axis: time across session. Y-axis: depth along electrode shank. Points (white marks) indicate individual spikes. Banding pattern reflects degree of motion before (left) and after (right) motion correction.

Based on coregistration between intraoperative navigation and preoperative imaging, postoperative high-resolution CT, and electrophysiological properties, we expect our units to be drawn mostly from CA1 with some possible recording sites in DG at the distal end (**Figure 1B-D**). **Figure 1C** shows the probe in the tissue, imaged in one patient (patient 2). Overall, the quality of isolation was high, consistent with past studies using Neuropixels probes in humans. Typical example waveforms are shown in **Figure 1E**.

Participants listened to audio podcasts derived from publicly available sources (see **Methods**) while undergoing surgical procedures. We performed recordings under general anesthesia; our previous work shows that many features of semantic encoding are preserved under general anesthesia (Katlowitz et al., 2025A). We extracted embeddings at the word level and sentence level from GPT-2, Word2Vec, and SBERT (see **Methods**). We used a standard analysis platform developed in our laboratory to quantify neural responses to speech (Franch et al., 2025; Katlowitz et al. 2025A and B; **Figure 1F**). Specifically, speech was carefully transcribed manually using Praat (Boersma & Weenink, 2025) by laboratory members and checked by the first author (EAM) so that word onset and offset times could be reconciled with neural responses at high resolution. Average firing rates were 1.6 +/- 1.2 Hz. Motion artifacts, a major challenge for human cortical Neuropixels recordings, were markedly less conspicuous in hippocampal recordings than in the cortical recordings (e.g., Paulk et al., 2022; **Figure 1G**).

### Semantotopy based on thresholded responses

To gain insight into the topography of semantic coding, we compared semantic tuning functions to each other, as a function of distance along the electrode shank. Specifically, we examined tuning function difference between all pairs of contacts, as a function of distance along the mediolateral axis. Thus, we first took all pairs of contacts located 20 microns apart (the shortest distance between two contacts along the shank) and averaged their semantic similarity (black arrows in **Figure 2A**). Then we took all cases in which contacts were located 40 microns away (blue lines in **Figure 2A**). We repeated this process with successively greater distances, up to a total distance of 3,060 microns (we chose this distance, which represents about 75% of the length of the active part of the probe, because, after that, the quantity of data gets too low to perform valid statistics).

**Figure 2.**
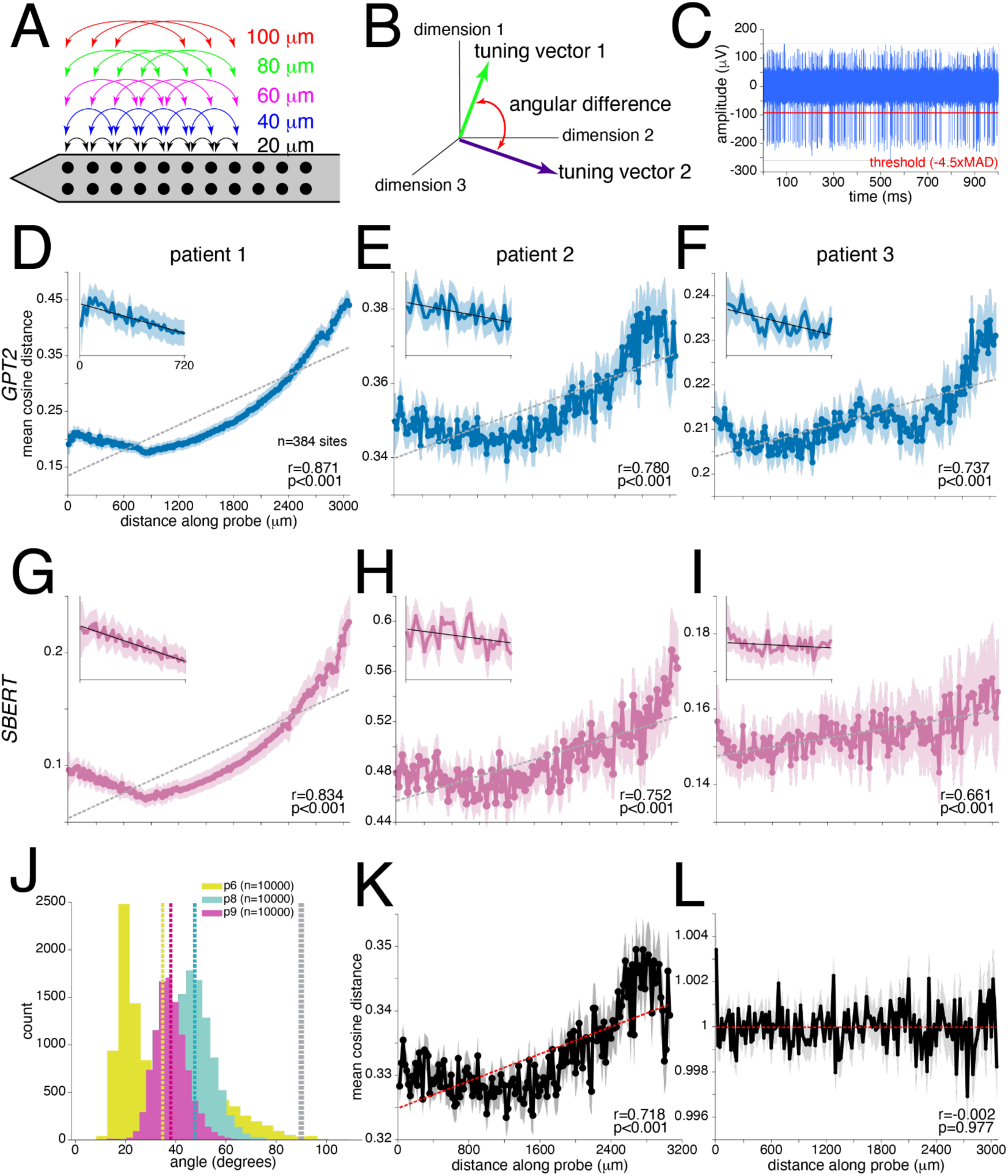
Semantic encoding along mediolateral axis of hippocampus with thresholded data. **A.** Schematic of distance analysis, illustrating which contacts were compared at distances along the probe. **B.** Schematic of angular difference calculation between pairs of tuning vectors, calculated from the predictor weights in the regression. **C.** Schematic of thresholding method. **D-F.** Cosine distance results as a function of probe distance from using GPT-2 embeddings in patient 1(D), patient 2 (E), and patient 3 (F). Inset: cosine distance results for the first 720 microns using GPT-2 embeddings. **G-I.** Cosine distance results as a function of probe distance from using SBERT embeddings in patient 1 (G), patient 2 (H), and patient 3 (I). Inset: cosine distance results for the first 720 microns using SBERT embeddings. **J.** Distributions of angular distance between 10,000 pairs of neurons. Yellow (patient 1), blue (patient 2), and pink (patient 3) lines indicate and mean angular distance. The gray line indicates 90 degrees, expected for completely random neurons. **K.** Cosine distance results as a function of probe distance using neural activity across two halves of data in patient 2 using GPT-2 embeddings (patients 1 and 3 give similar results, data not shown). **L.** Cosine distance results as a function of probe distance using tuning functions calculated from the regression’s null model in patient 2 (patients 1 and 3 give similar results, data not shown). Shaded regions represent +/- SEM.

We regressed neural responses against the semantic embedding derived from GPT-2 (**Figure 2B**). We used layer 36, because our past studies (Franch et al., 2025) suggest that neural decoding reliably resembles embeddings in that layer. (We obtained similar results with layer 24; see below). We predicted each contact’s response count for a given word from the dimensionality-reduced embeddings, word duration, and their interactions using a Poisson linear model (see **Methods** and Franch et al., 2025). Note that for this analysis, we used threshold crossing events (**Figure 2C**). We set spike detection thresholds adaptively for each channel to meet a standardized detection rate (see **Methods**). For each channel, we estimated the noise level using the median absolute deviation (MAD) method. Then, we defined a thresholded event as T = k x MAD, where k was optimized through an iterative binary search algorithm (range: -15 to -2.5 MAD) to achieve the target spike rate within 1 Hz tolerance. Threshold crossing allows for a direct comparison across all contacts, and likely involves a mix of local units. Data showing similar results from isolated units are found in **Figure 3**.

**Figure 3.**
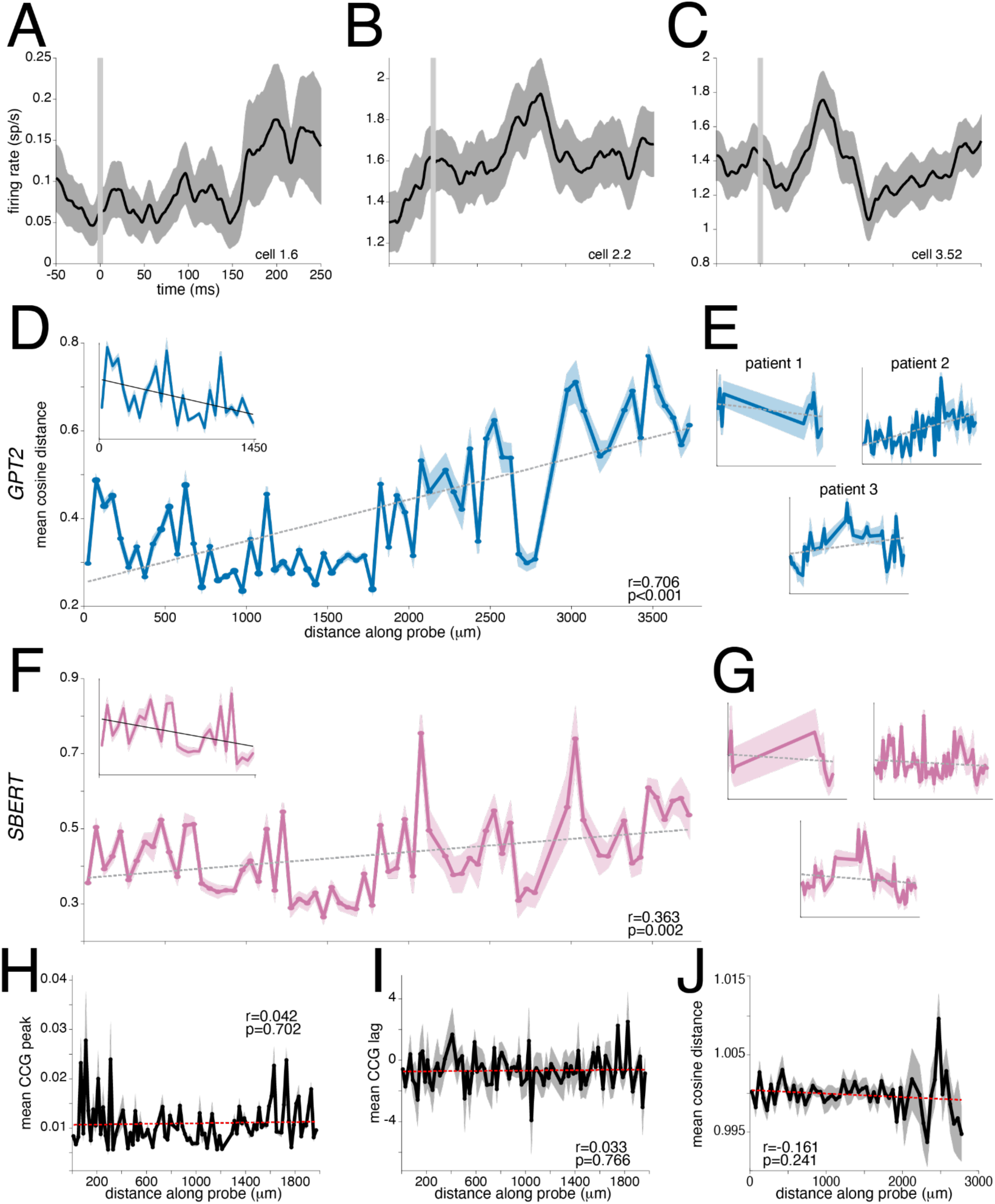
Semantic encoding along mediolateral axis of hippocampus with isolated units. **A.** Peri-stimulus time histogram demonstrating response to word onset from example neuron in patient 1. **B.** Peri-stimulus time histogram from example neuron in patient 2. **C.** Peri-stimulus time histogram from example neuron in patient 3. **D.** Cosine distance as a function of distance combined across patients using GPT-2 embeddings.. Inset: cosine distance as a function of distance combined across patients for shorter distances. **E.** Individual patient’s cosine distance as a function of distance using GPT-2 embeddings. **F.** Cosine distance as a function of distance combined across patients using SBERT embeddings.. Inset: cosine distance as a function of distance combined across patients for shorter distances. **G.** Individual patient’s cosine distance as a function of distance using SBERT embeddings. **H.** Mean CCG peak as a function of distance for patient 2. **I.** Mean CCG lag as a function of distance for patient 2. **J.** Cosine distance as a function of probe distance for patient 2 using tuning functions from the null model. Shaded regions represent +/- SEM.

Overall, we found a clear decrease in semantic similarity as a function of physical distance along the probe (Figure **2D****, E,** and **F**). Specifically, we observed a positive correlation between distance along the electrode shank and tuning curve cosine distance (patient 1: r = 0.871, p < 0.001; patient 2: r = 0.780, p < 0.001; patient 3: r = 0.737, p < 0.001). In other words, the tuning function at any recording site was more similar to tuning functions of closer sites.

GPT-2 is a contextual embedder; we also replicated the analysis with a non-contextual embedder, Word2vec (Mikolov et al., 2013), and found qualitatively identical results (patient 1: r = 0.897, p < 0.001; patient 2: r = 0.845, p < 0.001; patient 3: r = 0.835, p < 0.001; data not shown). Note that these results are based on embeddings derived from layer 36 in GPT-2. Some studies indicate that middle-to-late layers provide better match to neural data than the highest layer (Caucheteux et al., 2023). In this case, we do not find a large difference between them. Indeed, embeddings from layer 24 yielded similar results to layer 36, showing a significant decrease in semantic similarity as distance increases (patient 1: r = 0.906, p < 0.001; patient 2: r = 0.777, p < 0.001; patient 3: r = 0.774, p < 0.001, data not shown).

We also found a complementary effect of local semantic similarity at a smaller scale (**Figure 2D, E, and F insets**). Specifically, we found a small, but statistically significant, increase in semantic similarity as a function of distance, although only at short distances (∼600 to 800 microns). This effect was significant in all three patients (patient 1: r = -0.828, p < 0.001; patient 2: r = -0.553, p < 0.001; patient 3: r = -0.670, p < 0.001). This result is consistent with a periodic organization of semantic tuning, with a period of ∼700 microns. Note that this modest decrease cannot be explained by drift/uncorrected motion, as these would cause a positive, rather than negative correlation. It also cannot be explained by sampling the same neuron across multiple contacts, as semantic similarity would stay the same across multiple distances tested.

Thus, this negative slope would flatten and become weaker, demonstrating loss of effect size. Indeed, the modest decrease we observe is evidence that the large increase we observe is not due to motion correction artifacts, because those would cause the largest effects at the shortest distances. Finally, this effect is not likely to be driven by use of layer 36; using layer 24, we also found a significant increase in semantic similarity as distance increases for these shorter distances (patient 1: r = -0.726, p < 0.001; patient 2: r = -0.619, p < 0.001; patient 3: = -0.547, p < 0.001).

Meaning is carried at multiple levels in language (Gwilliams, 2025). Recent years have seen growing interest in sentence level embeddings, which can be inferred from a specialized embedder, SBERT. We next repeated the word-by-word analysis at the sentence level using SBERT regressions (**Figure 2G, H, and I**). We found largely similar results as word level.

Specifically, we found a gradual decrease in semantic similarity between sentence level semantic embeddings as a function of distance (patient 1: r = 0.834, p < 0.001; patient 2: r = 0.752, p < 0.001; patient 3: r = 0.661, p < 0.001). Notably, the local semantic similarity effect we observed in word level embeddings was also observed with sentence level embeddings (patient 1: r = -0.941, p < 0.001; patient 2: r = -0.357, p = 0.030; patient 3: r = -0.224, p = 0.183, **Figure 2G, H, and I insets**).

We performed a few control analyses to test whether these results could be attributed to low-level factors that would cause spurious results. Most importantly, we asked whether results were due to factors that reflect common data, such as possible model overfitting or common noise across channels. To control for these possible confounds, it is critical to use entirely unrelated data (both on the stimulus side and on the neural activity side) to fit semantic tuning curves. Thus, we performed split-half control, in which we estimated tuning curves using half the data and compared their semantic similarity with tuning curves estimates on the other half (using the regression approach described above). We computed 10 iterations of different halves. We found that angular difference between GPT-2 tuning vectors of random pairs of neurons across halves, computed for 10,000 pairs of neurons, was less than 90 degrees (patient 1: mean = 34.81 degrees; patient 2: mean = 47.58 degrees; patient 3: mean = 38.02 degrees, **Figure 2J**) but higher than between pairs of neurons trained and tested on the same half of data (patient 1: mean = 14.34 degrees; patient 2: mean = 17.55 degrees; patient 3: mean = 16.75 degrees, data not shown). The fact that the angles between random neurons across the two halves is less than 90 degrees indicates that results are due to semantic relationships between neurons rather than noise. Critically, the we observed the pattern of a decrease in semantic similarity as a function of distance for neurons across halves (patient 1: r = 0.871, p < 0.001; patient 2: r = 0.718, p < 0.001, patient 3: r = 0.678, p < 0.001; patient 2 illustrated in **Figure 2K**). Thus, the observed semantic and spatial relationships in **Figure 2D-I** are not due to noise.

Conversely, we were concerned that our results may reflect some inherent noise properties of the neurons such that, for example, perhaps adjacent sites have more similar intrinsic (that is, unrelated to the stimulus) neural firing rate patterns, and these somehow influence the estimated tuning curves. We therefore performed an analysis in which we calculated tuning vectors from weights in the null model where embeddings have been shuffled, but neural activity is not changed. This control breaks the relationship between neural activity and semantics, but keeps any intrinsic firing rate patterns. We no longer find a significant relationship between semantics and anatomical distance when using the null model (patient 1: r = -0.049, p = 0.543; patient 2: r = -0.002, p = 0.977; patient 3: r = -0.033, p = 0.686, patient 2 is shown in **Figure 2L**). This result indicates that the patterns we observe are not due to intrinsic properties of neural firing.

### Semantotopy based on unit responses

To ensure that the effects we observed were not somehow due to something about the thresholding process, we next confirmed that these effects can also be observed using isolated units (**Figure 3**). Neuropixel electrodes do not, as a rule, produce isolated units at each contact point. In our case we obtained a total of 23, 172, and 93 units in patients 1, 2, and 3, respectively. Responses of three example units are shown in **Figure 3A, B, and C**. These three units illustrate a typical dynamic range of responses and show typical temporal envelopes of word-evoked responses in our dataset.

We find that both patterns reported above (semantotopy at two spatial scales) are replicated using units, although, because of the increased noise, they are only observed when we combine across patients. Specifically, we found an overall decrease in semantic similarity as a function of unit distance across the group of three participants (r = 0.706, p < 0.001, **Figure 3D**). Individually, we do found statistical significance for patients 2 and 3 (patient 2: r = 0.555, p < 0.001; patient 3: r = 0.310, p = 0.048) but not patient 1 (who had the lowest number of single units; r = -0.352, p = 0.289, **Figure 3E**). We also found the same local semantic similarity effect where there is a significant increase in semantic similarity as a function of distance for short distances (r = -0.447, p = 0.015, **Figure 3D inset**); this effect was not replicated in any of the patients individually (p > 0.05 in all cases).

With word2vec, we also replicated the finding of a decrease in semantic similarity with distance (r = 0.0643, p < 0.001, data not shown), while the other effect, the local semantic similarity effect, approached significance (r = -0.365, p = 0.052, data not shown). Likewise, we found an overall decrease in semantic similarity using SBERT as a function of distance (r = 0.363, p = 0.002, **Figure 3F and G**). And, again for the SBERT analysis, we found the local semantic similarity effect as a function of distance (r = -0.396, p = 0.033, **Figure 3F inset**).

We performed the same control analyses on isolated units too. Moreover, because we were using single units, we tested functional connectivity, reasoning that sites might show strong functional connectivity with adjacent sites, and that these functional connections could spuriously cause correlations in tuning functions. Overall, we found no effect on the CCG peak, the peak in a cross-correlogram between two neurons, as a function of distance either at the level of individual patients or in the group (patient 1: r = 0.256, p = 0.579; patient 2: r = 0.042, p = 0.702; patient 3: r = 0.207, p = 0.153, patient 2 illustrated in **Figure 2H**). We also tested whether timing differences between neurons as a function of distance may indicate signs of functional connectivity. However, we did not find an effect on the CCG lag as a function of distance (patient 1: r = 0.574, p = 0.178; patient 2: r = 0.033, p = 0.766; patient 3: r = -0.049, p = 0.737, patient 2 illustrated in **Figure 2I**). Additionally, we tested these results with tuning functions from the regression’s null model. We found no significant results of cosine distance as a function of distance (patient 1: r = 0.836, p 0.370, patient 2: r = 0.161, p = 0.241, patient 3: r = -0.031, p = 0.845 patient 2 illustrated in **Figure 2J**). Lastly, we performed the same split-half analysis and found that the main result of semantic similarity decreasing as a function of distance was significant in patient 2 (r = 0.742, p < 0.001) and patient 3 (r = 0.510, p < 0.001) but not patient 1 (r = -0.008, p = 0.989, fewest amount of neurons, data not shown).

### Changes in semantic encoding along the mediolateral axis

We computed the best fit line for each of the 30 elements in the PCA-compressed tuning vector along the mediolateral axis to improve our estimates of the tuning along the shank (**Figure 4A**). We then computed the differential tuning function by subtracting each end’s tuning from its complement on the other end, and the centrer from both ends. We then ran 30-PCA reduced semantic embeddings from every word in a large database of 370,079 words (Ward, 2002) through estimated tuning functions for the medial and lateral ends of the probe, as well as the center. (We used Word2Vec for this analysis because words here were not contexualized by surrounding language). This analysis gives the words that are most differentially likely to drive responding at each recording site relative to the others.

**Figure 4.**
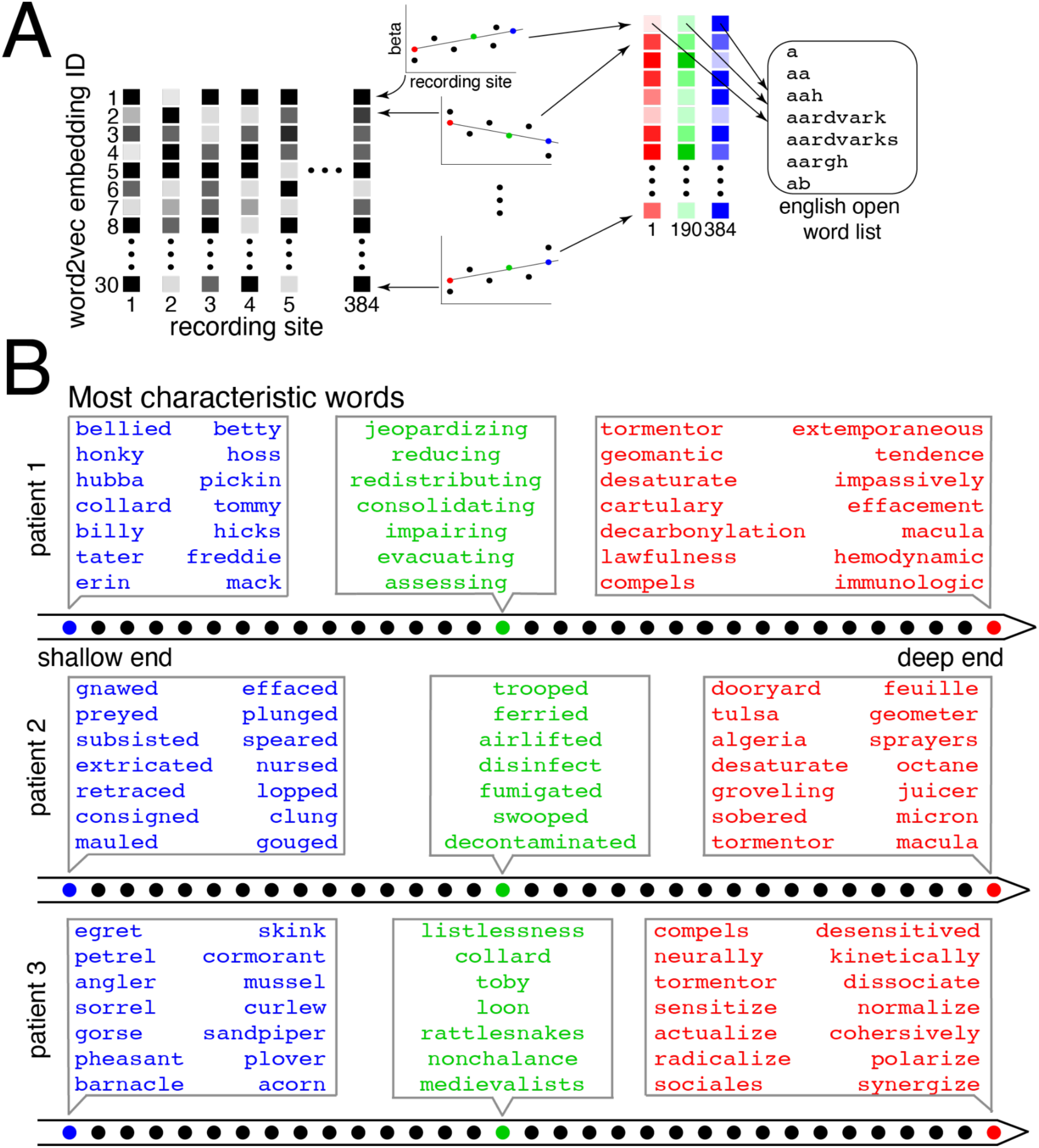
Words most characteristic of contacts along the mediolateral axis of hippocampus. **A.** Schematic of method used to calculate words in panel B. We used the same regression methods in Figure 2 **and 3** with word2vec embeddings. We fit lines to the tuning vectors for the 30 embedding dimensions across all neurons. These lines of best fit predicted responses at the most lateral, central, and medial contacts. We ran these lines of best fit against embeddings from a large corpus to determine words that are characteristic of lateral, central, and medial contacts. **B.** Words that are most differentially effective at driving neural activity in the lateral (blue), central (green), and medial (red) portions of our recording. These are words that were not tested during the passive listening experiment, but were derived from a large corpus, and that are most effective at driving fit tuning curves in each of three locations relative to the other two. Preferred words are characteristic within patient, but not consistent across patients.

For patient 1, the words at the proximal end of the probe (and thus, more laterally positioned on the head of the hippocampus), involve a greater degree of regional and slang terms, while words most characteristic of the deeper end of the probe (more medial within the hippocampus) are more associated with academia, scholarship, the legal system (**Figure 4B**). In the middle of the probe, the most characteristic words are formal and process-oriented. For patient 2, the lateral end is dominated by past-tense verb forms, while the medial end does not show an obvious pattern. Words located centrally are verbs related to transportation and cleaning. For patient 3, the lateral end is dominated by words that come from nature, while the medial end is dominated by present tense verbs. Centrally, there is no clear pattern. These results suggest that there are some semantic gradients across the mediolateral axis of the hippocampus, possibly several, and they are not consistent across individual patients.

To quantify these descriptive results, we regressed each of several semantic variables against position along the probe (**Figure 5**). To do this, we calculated a change vector for the same tuning vectors in **Figure 4** of each contact using least-squares regression. Then for each semantic variable we averaged the embeddings of most characteristic words subtracted by the average of the embeddings of the least characteristic words (see **Methods** for details). We then projected the semantic vector onto the change vector. We calculated correlations between responses to the semantic vector as a function of distance as shown in **Figure 5**. Lastly, we ran permutation tests (10,000 permutations) before calculating a two-tailed t-test between the two vectors.

**Figure 5.**
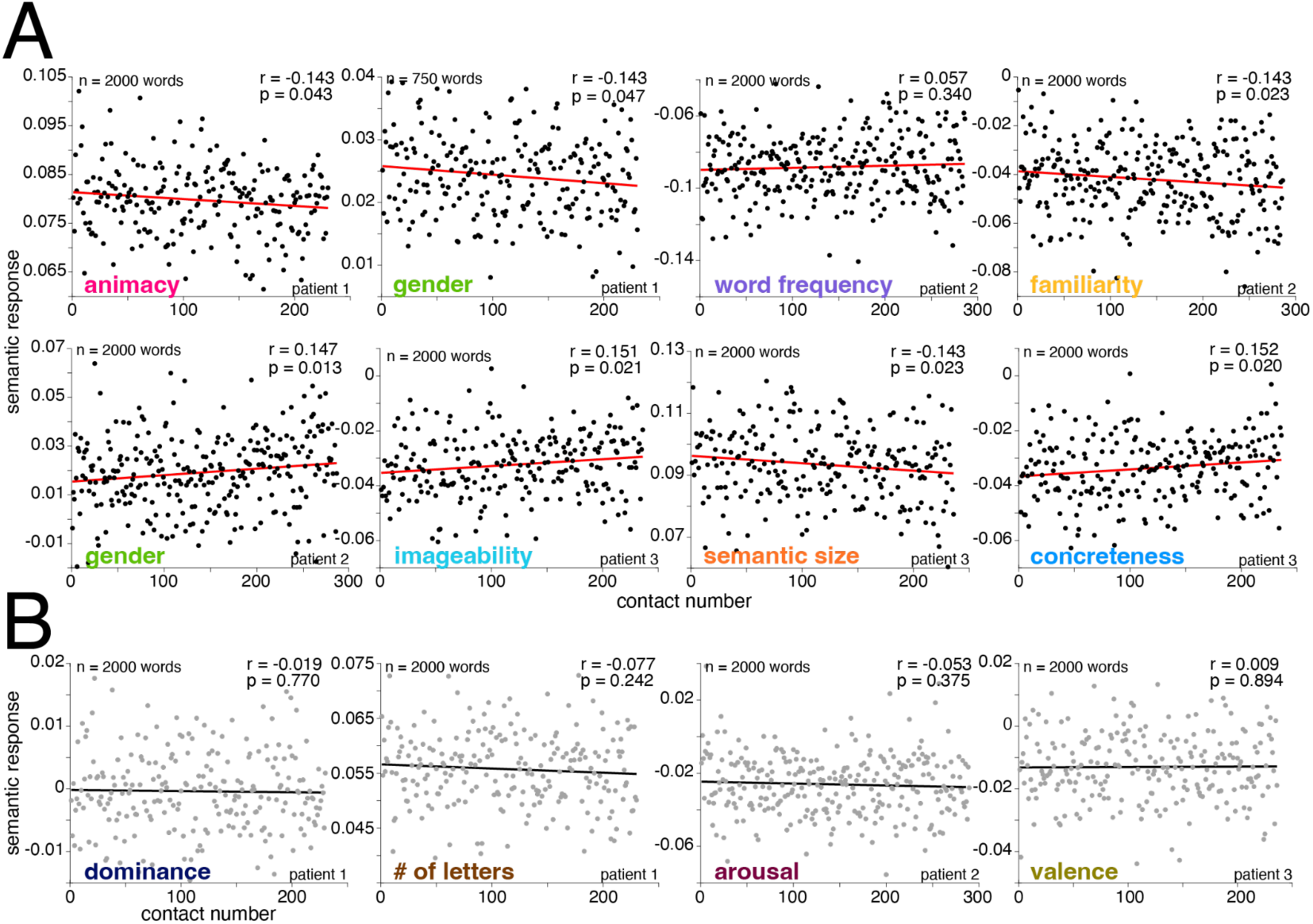
Semantic effects along mediolateral axis of hippocampus. **A.** Correlation of semantic response variable and distance along the probe. Semantic variables demonstrating significant responses as a function of depth included: animacy, familiarity, word frequency, concreteness, imageability, semantic size, and gender. **B.** Semantic variables that are not significantly correlated with function of depth include dominance, number of letters, arousal, and valence.

Some of the semantic variables showed significant changes with distance. Seven of the 13 variables tested were significant in at least one patient (this proportion is significantly greater than the false positive rate expected by chance, p=0.002; binomial test). In patient 1, animacy (Astbury, 2023) decreases from medial to lateral positioning within the hippocampus, p = 0.036, permutation test), meaning more medial contacts prefer more animate words. More medial sites also preferred more gendered words (Scott et al., 2019) in patient 1 (p = 0.047). In patient 2, more medial sites preferred less frequent words (Brysbaert et al., 2009), more familiar words (Scott et al., 2019), and less gendered words (Scott et al., 2019, p < 0.001; p = 0.031; p = 0.016). In patient 3, medial contacts also preferred less imaginable words (Scott et al., 2019), words greater in semantic size (e.g., airplane vs needle, Scott et al., 2019), and less concrete words (Brysbaert et al., 2014, p = 0.027; p = 0.035; p = 0.019). Tested variables that did not show any effect in any patient were number of letters, dominance, arousal, valence (Warriner et al., 2013, p > 0.05 in all cases), age of acquisition, and number of syllables (p > 0.05 in all cases, data not shown).

## DISCUSSION

We investigated the mapping of semantic information in the human hippocampus. We analyzed a rare dataset consisting of neural responses along the shank of a Neuropixels electrode placed mediolaterally within the head of the hippocampus during passive language listening. We found two distinct patterns at different scales. The larger effect was a gradual decrease in tuning similarity as a function of distance between pairs of recording sites at the scale of several millimeters. The second effect, the local semantic similarity effect, was an increase in semantic similarity as a function of distance. This effect was weaker and at a shorter scale (∼700 microns), where recording sites were most similar with those located a fixed distance away. This effect was also present for both word and sentence level embeddings in thresholded data and isolated units. These results are consistent with a periodic organization (that is, of repeating similarity at a fixed distance) of semantic tuning in the hippocampus. Together, these two effects indicate that hippocampal neurons are organized *semantotopically*, so that neurons with semantically similar tuning are close to each other along the electrode.

The broader semantotopy is reflected in differences in tuning within patients in the words themselves that drive a maximal response in contacts at different distances as well as the types of word (**Figure 4** and **Figure 5**). These findings suggest that not only is there an organization at the word level but also some general trends exist as gradients along the mediolateral axis. Given the variability in patients’ semantic topography shown in Figure 4 and Figure 5, future studies may want to examine to what extent language and life experience shapes these individual maps. For example, the regional slang that is maximally encoded in lateral regions of the hippocampus in patient 1 may be reflective of a specific cultural background, while the words related to nature maximally encoded in the lateral region in patient 3 may reflect a specific hobby. Future studies could perform similar analyses across patients of many different ages and backgrounds to see if semantic maps are more similar across patients at a young age and then become individualized with increased experience.

Why would the brain have a semantotopic map? Neurons with similar semantic tuning may cluster spatially in the hippocampus because of converging anatomical, computational, and developmental factors. Topographically organized inputs from the entorhinal and perirhinal cortex can deliver overlapping information about related semantic categories to neighboring hippocampal neurons, while short-range recurrent and inhibitory connections promote Hebbian clustering of co-active units that encode related concepts. Such local organization may also reflect wiring efficiency: grouping neurons with similar tuning minimizes axonal length, metabolic cost, and conduction delays (Laughlin & Sejnowski, 2003; Chklovskii & Koulakov, 2004). Once established, this proximity facilitates pattern completion and semantic generalization—allowing related memories or concepts (e.g., “dog,” “cat,” “pet”) to mutually reinforce one another during recall. Developmentally, gradients of molecular guidance cues and shared neurogenesis timing may bias nearby neurons toward similar input patterns, predisposing them to encode related semantic dimensions (Sur & Leamey, 2001; Donato et al., 2017).

Alternatively, these effects may be the result of learning-related processes throughout life (McClelland et al., 1995; Kumaran et al., 2016). Together, these mechanisms suggest that the hippocampus—despite its role in orthogonalizing distinct episodes—also supports local pockets of semantic coherence, where neighboring neurons represent conceptually related content through shared input structure, co-activity–driven plasticity, and energy-efficient network design.

The apparent periodic organization we observed could be a result of several factors. First, as hypothesized by Quiroga and others, it may be that bringing together dissimilar information makes conjunctive coding more efficient (Quiroga et al., 2005; Singer & Gray, 1995; Erez et al., 2016). One reason may be employment of a contrastive coding scheme in the hippocampus. It could be the case that neighboring neurons while similar are not as similar to neurons just a bit farther away in order to reduce redundancy. It would be efficient for neighboring neurons to carry similar though not identical information. This periodic organization may also indicate functional organization in the hippocampus. We may be recording from layers in the hippocampus where neighboring neurons are not functionally connected but those a bit farther away are. In this case, information may be more shared with neurons that are close but not neighboring. Both options would demonstrate this periodic organization and support efficient coding in the hippocampus.

This study has several limitations. First, this study focuses on a small part of the hippocampus and only the mediolateral axis. Future studies should examine how results reported here compare to those that are in different areas of the hippocampus or along other axes. (In particular, there is growing evidence that the anterior-posterior axis may have important featural differences, Moser & Moser, 1998; Poppenk et al., 2013; Strange et al., 2014). In addition, this study includes only three patients; results may differ with more patients being tested. Moreover, results are from anesthetized patients. While we have shown that semantic processing occurs during anesthesia, the organization of such representations may differ in patients who are awake during language comprehension tasks.

One particularly important reason for understanding the semantotopic maps relates to the importance of semantic encoding for brain computer interfaces (BCIs). Current approaches to BCI treat anarthrias and dysarthrias, not aphasias. However, BCIs that can read out semantic information can potentially treat aphasia. In order to do semantic readout, we need to understand the basis of semantic encoding. Likewise, understanding specific organization of concepts in the hippocampus may contribute to stimulation techniques that aim to restore memory (Roeder et al., 2022; Bick and Eskandar, 2016). Furthermore, these principles of organization may also be applied to speech prosthesis. While speech neuroprosthesis already targets the hippocampus along with other regions, results focus on articulatory features of language (Stavisky, 2025). Including semantic concepts may only improve performance. Knowing the organization of concepts in the hippocampus can potentially help with the design of recording technologies that assist in converting thoughts to synthetic speech.

## METHODS

### Patient recruitment

Experiments were conducted according to protocol guidelines approved by the Institutional Review Board for Baylor College of Medicine, Houston TX (H-50885). All recruited patients were diagnosed with drug resistant temporal lobe epilepsy and were scheduled to undergo an anteromesial temporal lobectomy for seizure control. All patients provided written informed consent to participate in the study and were aware that participation was voluntary and would not affect their clinical course. Included patients’ age ranged from 25-46 years old (25, 45, and 46 years old), with two males and one female. Patients 1 and 3 were native English speakers; patient 2 was a bilingual English and Spanish speaker. Two resections were on the left side, and one was on the right. All patients were right handed. None of the patients reported explicit memory of intraoperative events after the case when discussed in the post-operative care unit or while recovering in the hospital the next day.

### Experimental stimuli

Patient 1 listened to three stories from *The Moth Radio Hour*, totaling 19:28 minutes and 3,024 words. The three stories were, “Juggling and Jesus,” “Wild Women and Dancing Queens,” and “My Father’s Hands.” In each story, a single speaker tells an autobiographical narrative in front of a live audience. Patients 2 and 3 listened to “Why We Should NOT Look for Aliens - The Dark Forest,” an educational video created by the Kurzgesagt group (Kurzgesagt GmbH; Munich, Germany). These story totaled 10:31 minutes and 1,565 words.

### Neuropixels data acquisition setup and intraoperative recordings

Neuropixels 1.0-S probes (IMEC) with 384 recording channels (total recording contacts = 960, usable recording contacts = 384) were used for recordings (dimensions: 70µm width, 100µm thickness, 10mm length). The Neuropixels probe, consisting of both the recording shank and the headstage, were individually sterilized with ethylene oxide (Bioseal, CA, Coughlin 2023). Our intraoperative data acquisition system included a custom-built rig including a PXI chassis affixed with an IMEC/Neuropixels PXIe Acquisition module (PXIe-1071) and National Instruments DAQ (PXI466 6133) for acquiring neuronal signals and any other task-relevant analog/digital signals respectively. Our recording rig was certified by the Biomedical Engineering at Baylor St. Luke’s Medical Center, where the intraoperative recording experiments were conducted. A high-performance computer (10-core processor) was used for neural data acquisition using open-source software such as SpikeGLX 3.0 and OpenEphys version 0.6x for data acquisition (AP band (spiking data)), band-pass filtered from 0.3kHz to 10kHz was acquired at 30kHz sampling rate; LFP band, band-pass filtered from 0.5Hz to 500Hz, was acquired at 2500Hz sampling rate). We used a “short-map” probe channel configuration for recording, selecting the 384 contacts located along the bottom 1/3 of the recording shank.

Audio was played via a separate computer using pre-generated wav files and captured at 30kHz or 1,000kHz on the NIDAQ via a coaxial cable splitter that sent the same signal to speakers adjacent to the patient. MATLAB (MathWorks, Inc.; Natick, MA) in conjunction with a LabJack (LabJack U6; Lakewood, CO) was used to generate a continuous TTL pulse whose width was modulated by the current timestamp and recorded on both the neural and audio datafiles. Online synchronization of the AP and LFP files was performed by the OpenEphys recording software. Offline synchronization of the neural and audio data was performed by calculating a scale and offset factor via a linear regression between the time stamps of the reconstructed TTL pulses and confirmed with visual inspection of the aligned traces.

Acute intraoperative recordings were conducted in brain tissue designated for resection based on purely clinical considerations. The probe was positioned using a ROSA ONE Brain (Zimmer Biomet) robotic arm and lowered into the brain 5-6mm from the ependymal surface using an AlphaOmega microdrive. The penetration was monitored via online visualization of the neuronal data and through direct visualization with the operating microscope (Kinevo 900). Reference and ground signals on the Neuropixels probe were acquired separately by connecting to a sterile microneedle placed in the scalp (separate needles inserted at distinct scalp locations for ground and reference respectively).

For all patients (n=3), we conducted neuronal recordings under general anesthesia for at most 30 minutes as per the experimental protocol. All patients were under total intravenous anesthesia, with propofol as the main anesthetic per experimental protocol. Inhaled anesthetics were only used for induction and stopped at least an hour prior to recordings. The anesthesiologist titrated the anesthetic drug infusion rates so that the BIS monitor (Medtronic; Minneapolis, MN) value was between 45 and 60 for the duration of the surgical case (Singh 1999). Of note, BIS values range between 0 (completely comatose) and 100 (fully awake), with standard intraoperative values to be between 40 and 60. We then carried out hippocampal recordings after resection of the lateral temporal lobe but prior to any resection of the hippocampus.

### Micro CT

Since recordings were only performed in tissue planned for resection, we first removed a small cube of tissue around the probe and then proceeded with the remainder of the resection. The cube specimens were processed following previously described methods (Hsu et al., 2019). In brief, resected specimens were fixed in 4% PFA for 16 hours at 4°C. They were then stabilized using a modified Stability buffer (mStability), containing 4% acrylamide (BIO-RAD, cat. no. 1610140), 0.25% w/v VA044 (Wako Chemical, cat. no. 017-19362), 0.05% w/v saponin (MilliporeSigma, cat. no. 84510), and 0.1% sodium azide (MilliporeSigma, cat. no. S2002). Samples were equilibrated in the hydrogel solution for 16 hours at 4°C before undergoing thermo-induced crosslinking at -90kPa and 37°C for 3 hours. Following crosslinking, excess hydrogel solution was removed, and specimens were washed four times with 1X PBS. Next, samples were immersed in 0.1N iodine and incubated with gentle agitation for 24 hours at room temperature before being embedded in agarose and imaged using a Zeiss Xradia Context micro-CT at 3µm/ voxel resolution. The acquired back-projection images were reconstructed using Scout-and-Scan Reconstructor (Carl Zeiss, Ver. 16.8) and converted to NRRD format via Harwell Automated Recon Processor (HARP, Ver. 2.4.1, Brown et al., 2018), an open-source, cross-platform application developed in Python. The 3D volumes were analyzed, and optical sections were captured using 3D Slicer (Fedorov et al., 2012).

### Neuronal data processing

#### Motion correction

We utilized previously developed and validated motion estimation and interpolation algorithms to correct for the motion artifacts from brain movement (Windolf et al., 2025). Motion was estimated via the DREDge software package (Decentralized Registration of Electrophysiology Data software, https://github.com/evarol/DREDge) using either a combination of motion traces obtained using raw LFP and/or AP band data, fine-tuned for individual recordings. Motion-correction was then implemented using interpolation methods (https://github.com/williamunoz/InterpolationAfterDREDge). Both the AP and LFP band data are motion-corrected and utilized for further pre-processing and analysis steps. If the estimated motion led to no improvement in the spike locations then spike sorting proceeded with the motion correction package built into Kilosort 4 without performing interpolation.

#### Unit extraction and classification

For analyses utilizing isolated units (**Figure 3**), automated spike detection and clustering were performed by Kilosort 2.0 if motion correction was already applied using the DREDge algorithm or KiloSort 4.0 (Pachitariu et al., 2024) if motion correction was not applied separately. Manually curation of spike clustered was performed using the open-source software Phy (Rossant et al., 2016). Unit quality metrics were calculated using SpikeInterface (Buccino et al., 2020) and were considered single units if they had a d-prime (d’) greater than 1 and fewer than 3% of spikes were violations of a 2ms inter-spike interval refractory period.

### Motional Analysis

The motion-corrected location estimates were obtained at a 250Hz sampling frequency using the DREDge algorithm. This signal was downsampled to 10Hz. The power spectrum of the calculated motion was then estimated using Welch’s overlapped segment averaging estimator for frequencies between 0.1 and 3Hz. The amount of motion was defined as the root mean square error of the location trace of the probes center relative to its average location.

### Audio transcription

After experiments, the audio.wav file was transcribed using AssemblyAI, a state-of-the-art AI model trained to transcribe speech. The transcribed words and corresponding timestamps from AssemblyAI were converted to a .TextGrid and loaded into Praat (Boersma & Weenink, 2025), a well-established software for speech analysis, with the original audio.wav file. Trained lab members used the spectrogram and waveform to correct each word onset and offset. The .TextGrid output of corrected words and timestamps from Praat was converted to an Excel file and loaded into MATLAB for further analysis.

### Firing rate responses to words

The firing rate of each neuron for each word was the number of spikes that occurred during a word’s duration with an 80 ms delay. This value was divided by word duration and then multiplied by 1000 to compute spikes per second.

### Thresholding method (Figure 2)

Spike detection was performed using an adaptive thresholding algorithm designed to achieve a target firing rate of approximately 20 Hz per channel. A threshold was set dynamically for each channel based on signal characteristics and noise level. For each channel noise level was estimated using the mean absolute deviation (MAD), calculated as MAD = median (|x|)/0.6745, where x is the band-passed filtered signal. Channels with MAD values below 1x10^-6^ were classified as dead channels and excluded. An iterative binary search algorithm was used to determine the optimate threshold multiplier for each channel. Different values were tested within a range. Negative-going threshold crossings were considered a neuronal spike. If the iterative firing rate was too high, the threshold was made more negative and vice versa for a low firing rate. A refractory period of 1ms was also used to eliminate some level of noise.

### Regression spike counts on word and sentence embeddings

Word embeddings were calculated using the GPT-2 large model (Radford et al., 2019), a contextual word embedder. Layer 36 was used in all analyses, though results were similar using layer 24. Sentence embeddings were calculated using SBERT (Reimers & Gurevych, 2019). The last layer was also used for SBERT. We first used Principle Component Analysis on the full embeddings to obtain uncorrelated features that still capture the dominant structure in the embeddings space with reduced dimensionality. For each word, we used the first 30 principle components (PCs) from GPT-2 and 5 PCs for SBERT.

We modeled spike count responses of individual neurons using a Poisson Generalized Linear Model (GLM) with a log-link function and ridge (L2) regularization). The model aimed to predict the number of spikes a neuron fired in response to each word (summed spikes across word duration), using the top 30 or 5 PCs of the embedding vector, the duration of the word, and the interaction between each PC and the word’s duration as predictors (z-scored prior to model fitting). This resulted in a 61- or 11-dimensional feature vector for each word: 30 or 5 PC values, 1 duration value, and 30 or 5 PC×duration interaction terms. The Poisson GLM assumes that the spike count y*_i_* for word *i* is drawn from a Poisson distribution with mean λᵢ where:

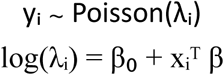

Where λ_ᵢ,*n*_, the expected spike count for word (*i*) and neuron n (*n*), is modeled by the linear regression coefficients (*β*), one of each of the 61 predictors (*x_i_^T^*), plus the y-intercept (*β*₀). For each neuron, we used a 10-iteration nested training loop, where in each iteration, the data was split into training and held-out test sets (80/20 split). Within the training set, 5-fold cross validation was used to select the optimal regularization strength (alpha) from a log-spaced range based on optimizing the cross-validated log-likelihood. The selected model was then evaluated on the held-out test set using several performance metrics, including Pearson correlation between predicted and observed spike counts, log-likelihood, and adjusted R².

Betas for each of the 30 or 5 PCs were used to calculate tuning vectors for each neuron. Pairs of tuning vectors were then related using cosine distance (1 - cosine angle). Distances with a minimum of 20 pairs of neurons were calculated. For the analyses with isolated data (Figure 3) bins of 50 microns were used.

This same analysis was performed with a range of neurons where firing rate was consistent to ensure increasing cosine distance as distance increases was not due to changes in firing rate in thresholded data.

The same method was used in a split-half version where instead of using the full set of data, neurons were trained and tested on two halves of data with 10 iterations. 10,000 pairs of neurons were computed for the distribution and mean angular distance.

### CCG analysis

Cross correlograms (CCGs) of semantic encoding were computed using a fixed duration of 300 ms following word onset by sliding the spike trains of each cell pair and counting coincident spikes within 1 ms time bins for each word and pair of neurons using the *xcorr* function in Matlab 2023b. Cross-correlations were normalized by the geometric mean spike rate to account for changes in individual neuron’s firing rate, and further corrected for stimulus-induced correlations by subtracting an all-way shuffle predictor - efficiently computed from the cross-correlation of the peri-stimulus time histograms (PSTHs, Bair et al., 2001; Pojoga et al., 2020; Franch et al., 2024). Specifically, trial (word) averaged cross correlation of the binary time series spike trains between neurons j and k was computed as:

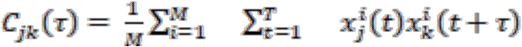

where M is the number of trials, T is the duration of the spike train segments, x is the neural response and τ is the lag. We normalized the above trial averaged cross correlation (Eq.1) by dividing it by triangle function and geometric mean of the average firing rates of the neurons 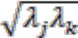 (Bair et al., 2001) to get the unbiased cross-correlogram of the spike trains in units of coincidences per spike:

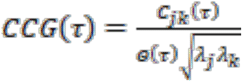

The function Θ(*τ*) is a triangle representing the extent of overlap of the spike trains as a function of the discrete time lag τ:

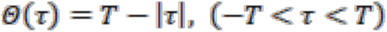

where T is the duration of the spike train segments used to compute *C_jk_* (Bair et al., 2001). Dividing *C_jk_* by Θ(τ) corrects for the triangular shape of *C_jk_* caused by the finite duration of the data (Bair et al., 2001). Dividing by 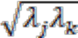 in Eq.2 results in CCG peaks with relatively constant area as firing rates of individual neurons change (Bair et al., 2001; Kruger & Aiple, 1988). In other words, dividing by the geometric mean of the firing rates of the two neurons makes the CCG peaks relatively independent of the firing rates. The maximum value, or peak, of each word’s CCG was calculated for all pairs of neurons at all distances (using the same distance calculations as the semantic similarity and distance calculations). The time lag at which the CCG peak occurred after word onset was also calculated for all pairs of neurons at all distances.

### Characteristic words along the probe

The same Poissin GLM was used with word2vec embeddings (Joulin et al., 2017), non-contextual word embeddings. Embeddings were reduced to the first 30 PCs. Betas from the regression for each neuron (30 dimensions) were extracted. A linear regression was used to estimate the slope and intercept of betas across all neurons for each of the 30 dimensions. This estimated line of best fit acts as a “pure” tuning vector for each neuron across the 30 dimensions. The array of lines of best fit were chosen for the deepest, shallowest, and middle contacts. Then a line of best fit was fit to tuning vectors for each embedding dimension across all neurons. Responses at these contacts were predicted from 30-dimensional PCA reduced word2vec embeddings of a large English corpus. Then, 60 words that were characteristic of the extreme ends of the probe and the middle were calculated.

### Semantic variables analyses

Ratings for different semantic variables were extracted from various databases. Semantic category vectors were created by subtracting the embeddings of the least characteristic words from the embeddings of the most characteristic words. Word2vec embeddings were used for all analyses. All analyses used the top 2,000 and bottom 2,000 words, except for the gender analysis with patient 1 which used the top 750 and bottom 750 words. A change vector was also calculated by performing least-squares regression on the tuning vectors from all 384 contacts. We then tested whether the semantic variable of interest aligned with the change vector by projecting the semantic vector onto the change vector, using the dot product. Permutation testing (10,000 permutations, shuffling word-semantic assignments) and bootstrap confidence intervals (10,000 iterations) were used to determine statistical significance.

## Acknowledgements

We thank the following people for valuable assistance: Joshua Adkinson, Garrett P. Banks, Sydney S. Cash, Victoria Gates, Chih-Wei Hsu, Domokos Meszéna, William Muñoz, Angelique C. Paulk, Andrew Watrous, Ziv Williams.

